# Transcranial alternating current stimulation with speech envelopes modulates speech comprehension

**DOI:** 10.1101/097576

**Authors:** Anna Wilsch, Toralf Neuling, Jonas Obleser, Christoph S. Herrmann

## Abstract

Cortical entrainment of the auditory cortex to the broadband temporal envelope of a speech signal is crucial for speech comprehension. Entrainment results in phases of high and low neural excitability, which structure and decode the incoming speech signal. Entrainment to speech is strongest in the theta frequency range (4–8 Hz), the average frequency of the speech envelope. If a speech signal is degraded, entrainment to the speech envelope is weaker and speech intelligibility declines. Besides perceptually evoked cortical entrainment, transcranial alternating current stimulation (tACS) entrains neural oscillations by applying an electric signal to the brain. Accordingly, tACS-induced entrainment in auditory cortex has been shown to improve auditory perception. The aim of the current study was to modulate speech intelligibility externally by means of tACS such that the electric current corresponds to the envelope of the presented speech stream (i.e., envelope-tACS). Participants performed the Oldenburg sentence test with sentences presented in noise in combination with envelope-tACS. Critically, tACS was induced at time lags of 0 to 250 ms in 50-ms steps relative to sentence onset (auditory stimuli were simultaneous to or preceded tACS). We performed single-subject sinusoidal, linear, and quadratic fits to the sentence comprehension performance across the time lags. We could show that the sinusoidal fit described the modulation of sentence comprehension best. Importantly, the average frequency of the sinusoidal fit was 5.12 Hz, corresponding to the peaks of the amplitude spectrum of the stimulated envelopes. This finding was supported by a significant 5-Hz peak in the average power spectrum of individual performance time series. Altogether, envelope tACS modulates intelligibility of speech in noise, presumably by enhancing and disrupting (time lag with in-or out-of-phase stimulation, respectively) cortical entrainment to the speech envelope in auditory cortex.

## 1. Introduction

Neural oscillations entrain to the temporal regularity of external stimuli. That is, a rhythmically presented stimulus leads to a phase and amplitude alignment of neural oscillations within the frequency of the presented rhythm (e.g., Thut et al., 2011). When listening to speech, auditory cortex tracks the spectro-temporal structure of speech (Abrams et al., 2008; Giraud and Poeppel, 2012; Lalor and Foxe, 2010; Luo and Poeppel, 2007). The strongest response can be found in the theta range around 4–8 Hz which corresponds to the average frequency rate of a speech signal (see Kayser et al., 2015, for a nuanced perspective of cortical responses to speech). Specifically, it reflects the broadband temporal envelope of the speech signal (Chandrasekaran et al., 2009; Ghitza and Greenberg, 2009). This kind of cortical entrainment to the envelope has been shown to be crucial for speech comprehension (e.g., Doelling et al., 2014). The functional role of cortical entrainment to the envelope has been widely discussed (for a review, see Ding & Simon, 2014;). Whereas some argue that entrainment to the envelope only tracks the acoustic properties of the envelope (Steinschneider et al., 2013; Millman et al., 2015) others argue that entrainment is a mechanism of syllabic parsing (Giraud and Poeppel, 2012) or sensory selection, (i.e., segragating the speech stream from background noise; Schroeder & Lakatos, 2009). Kösem and van Wassenhove (2017) suggest in their opinion paper that entrainment in the theta range processes acoustic and phonological information of the signal. This conclusion is strengthened by their findings on bistable speech sequences (Kösem et al., 2016). They were able to demonstrate that low frequencies were rather driven by the speech stream in a bottom-up fashion. Instead, higher frequencies were rather affected by top-down control and their activity was more related to the conscious speech percept. In line with these findings, Peelle and Davis (2012) argue that neural oscillations in the 4–8-Hz range structure and decode the incoming acoustic stream.

The extent to which auditory cortex is able to track speech depends on the amount of spectro-temporal detail in the signal. For example, a degraded speech signal leads to weaker neural entrainment to the envelope than clear speech (Peelle & Davis, 2012). Consequently the decline in speech intelligibility is stronger when acoustic information is missing (Ahissar et al., 2001; Peelle et al., 2013). Ding and Simon (2013) manipulated the signal-to-noise ratio of continuous speech and stationary noise showing that entrainment to the envelope in the 4 to 8-Hz range decreases with increasing noise level. The question that we would like to address with this study is whether it is possible to enhance cortical entrainment to the envelope when the intelligibility of speech is low (e.g., due to noise). Would in turn speech intelligibility increase when the neural signal-to-noise ratio is increased? For example Lorenzi et al. (1999) showed that raising the low-frequency amplitude of a speech signal masked with noise to the power of 2 increased intelligibility. They explain that the enhancement of the otherwise masked low-frequency temporal modulations led to increased intelligibility. However, we here introduce a different technique to modulate ongoing brain oscillations: transcranial alternating current stimulation (tACS). tACS induces cortical entrainment in a frequency specific manner (Antal & Paulus, 2013 and Herrmann et al., 2013; Helfrich et al., 2014; Zaehle et al., 2010). By modulating the resting membrane potential of neurons via tACS, oscillatory activity is guided by external oscillations (Fröhlich and McCormick, 2010). At the same time, tACS disrupts entrainment by inducing anti-phasic stimulation to synchronized cortical activity (Helfrich et al., 2014a; Strüber et al., 2014).

tACS induced entrainment has been previously shown to impact auditory perception (Heimrath et al., 2016). For example, tACS in the alpha frequency range (10 Hz) sinusoidally modulated detection of pure tones in noise (Neuling et al., 2012). Similarly, detection of near-threshold auditory click trains was also modulated by 4-Hz tACS phase (Riecke et al., 2015a). Both studies show that tACS-entrainment modulates the perception of near-threshold auditory stimuli. Furthermore, two studies investigating the impact of tACS on speech stimuli showed that phoneme detection is modulated by 40-Hz tACS in younger and in hearing impaired adults (Rufener et al., 2016; Rufener et al., 2016).

Since cortical entrainment to the speech envelope is critical for speech comprehension, and cortical entrainment to tACS has been shown to improve auditory perception, we hypothesize that speech intelligibility can be modulated with tACS. An electric current in the shape of the speech envelope (i.e., envelope-tACS) will be stimulated, while participants listen to speech-in-noise. Envelope-tACS will simulate the local field potentials during the presentation of clear speech. Critically, the timing of sentence presentation and tACS onset needs to be considered, because to enhance entrainment, tACS and the auditory-cortex response to the envelope need to be phase-aligned. While many studies report a duration of around 100 ms until auditory cortex presents a phase reset to incoming auditory information (e.g., Baltzell et al., 2016), other studies report longer durations ( 152 to 208 ms; Aiken and Picton, 2008). Riecke et al., (2015b) have shown that the perceptual buildup of an auditory stream can be modulated with tACS in the frequency of the stream. They varied the phase-lag between stream presentation and tACS phase demonstrating that the optimal phase lag (i.e., fastest perceptual build up) varied across participants. Furthermore, perceptual build up varied sinusoidally along the six different phase-lags of a 4-Hz sinewave. In accordance with the findings of Riecke and colleagues, we will approach this variability by manipulating the exact time lag between sentence presentation and tACS from 0 to 250 ms. This time lag manipulation will also allow for the observation of aligned/in-phase entrainment as well as out-of-phase or disrupting entrainment possibly impeding sentence comprehension. Overall, we expect to show that envelope-tACS in auditory cortex modulates sentence comprehension and that sentence comprehension fluctuates across time lags. Specifically, due to the circular nature of neural oscillations and the sinusoidal-like shape of the speech envelope and the neural excitability fluctuates accordingly (Lakatos et al., 2005). Therefore, we assume that sentence comprehension fluctuates sinusoidally along the different time lags (see also Riecke et al., 2015b).

## 2. Methods

### 2.1. Participants

Nineteen healthy young adults (11 female, mean age = 23.5 years) without prior or current neurological or psychiatric disorders participated in the study. The experimental protocol was approved of by the ethics committee of the University of Oldenburg, and has been performed in accordance with the ethical standards laid down in the 1964 Declaration of Helsinki. All participants gave written informed consent before the beginning of the experiment.

### 2.2. Experimental procedure

Each participant completed the experiment in two sessions on two different days in order to assure total tACS duration to be kept below 20 minutes per day. Participants were seated upright in a recliner in a dimly lit room before the stimulation electrodes and headphones were attached. Afterwards, the tACS threshold was determined. To this end, we started the stimulation at 3 Hz for a duration of 5 seconds with an intensity of 400 μA and asked the participants to indicate whether they perceived a skin sensation or phosphene. The stimulation intensity was increased by steps of 100 μA until the participant indicated skin sensation or phosphene perception or an intensity of 1500 μA was reached. In case the participant already reported an adverse effect at 400 μA, the intensity was reduced to a start level of 100 μA. Stimulation intensity throughout the experiment corresponded then to the highest intensity that was not perceived by the participant. Then, the participant’s absolute threshold of hearing was determined for each ear (i.e., subjective hearing threshold). After that, the participant was familiarized with the speech test (Oldenburger sentence test, see below) followed by the actual experiment. During the experiment, the participant completed nine lists of the test that were randomly assigned to the six tACS conditions and three control conditions (Figure 1A), which took around 45 minutes.

**Figure 1.**
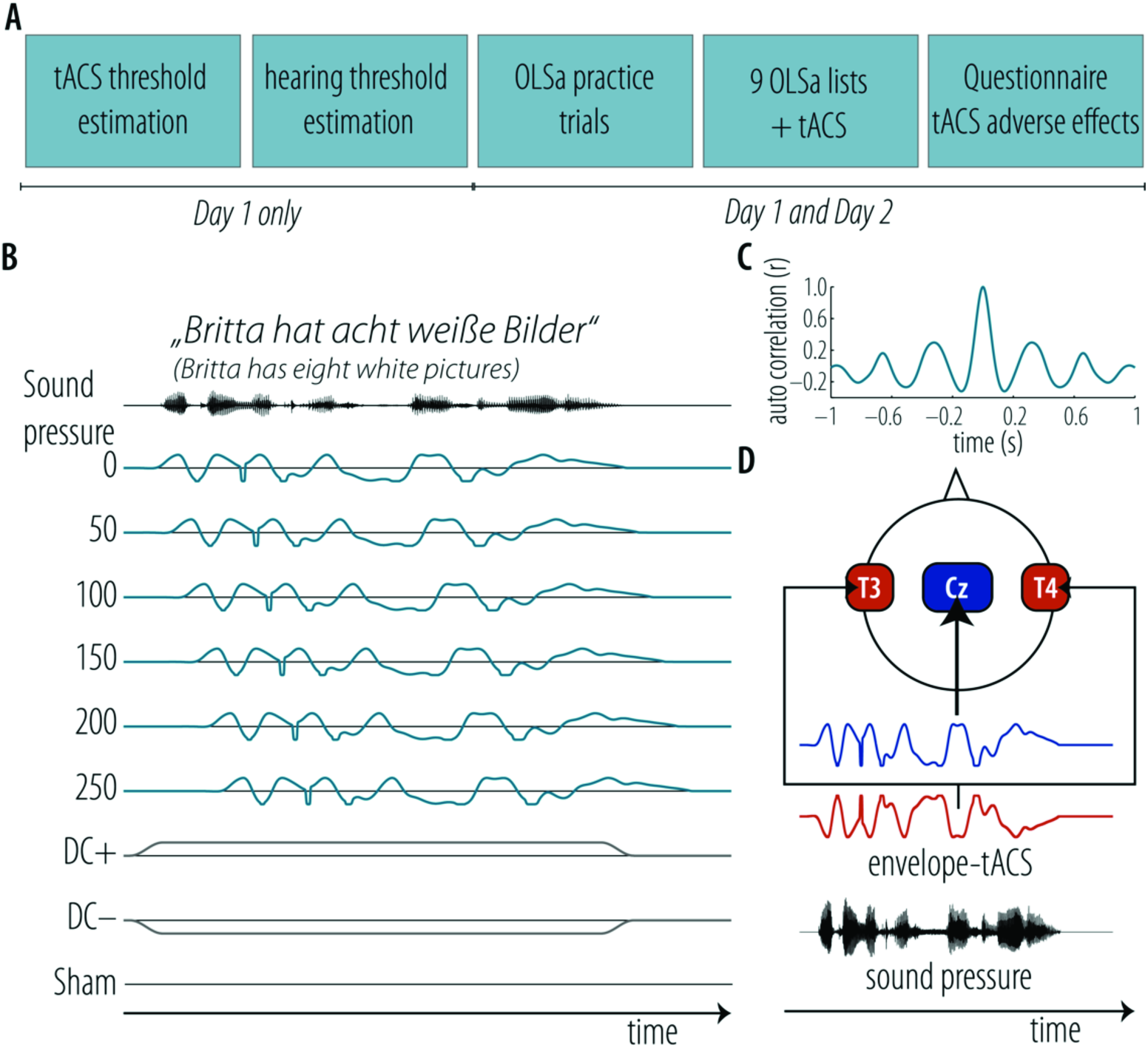
Experimental procedure, tACS conditions, and electrode configuration. **A.** Schematic display of the experimental time line. **B.** Example of a test sentence and the corresponding experimental conditions. Blue traces depict the six different time lags, gray traces the direct current conditions. Time lags refer to the point in time of tACS stimulation after presentation of the auditory sentence (i.e., 0–250 ms). **C**. Auto-correlation of one exemplary envelope (see also the corresponding average spectral power of all envelopes in Figure 4C). **D.** Blue and red boxes illustrate positions of the tACS electrodes on the head. Blue and red traces depict an example envelope of a sentence (black: audio signal) that was used for tACS stimulation.

The design was double blind, i.e., neither the participant, nor the experimenter knew the order of the conditions before the end of the experiment. After each list, the participants had to indicate whether they perceived the electrical stimulation. Following the speech test, the participants were asked to complete a questionnaire on possible adverse effects of tACS (Neuling et al., 2013). See Figure 1A for a schematic display of the experimental procedure.

#### 2.2.1. Oldenburg sentence test

The Oldenburg sentence test (OLSa; Wagner et al., 2001) is an adaptive test that can be used to obtain a speech comprehension threshold (SCT) during noise. Thirty sentences that consisted of five words are presented and the participant is asked to repeat the sentences orally. The test material consists of 40 test lists of which 22 (on two days) were randomly chosen for each participant. For each test session, the two test lists were used to familiarize the participant with the test material according to the handbook. Subsequently, nine lists were presented, separated by self-paced breaks. Note that due to the strong regularity of the 5-word sentences of the OLSa, the envelope fluctuations of these sentences are approximating a sinusoidal shaped. Therefore, they are specifically suited to investigate and modulate cortical entrainment to the speech envelope. Figure 1C illustrates this by plotting the autocorrelation of an exemplary sentence envelope.

The presented sentences were sampled at 44,100 Hz and delivered via MATLAB to a D/A converter (NIDAQ, National Instruments, TX, USA), attenuated separately for each ear (Tucker-Davis Technologies, Alachua, FL, USA; model PA5), and presented via headphones (HDA 200, Sennheiser, Germany). The noise masker of the OLSa was presented at 65 dB above individual hearing threshold; the intensity of the sentences was adjusted according to the test manual. The noise masker had the same spectral characteristics as the presented sentence, in order to obtain an optimal masking effect of the sentence. The noise masker was presented from 0.5 s prior to sentence onset until 0.5 s after sentence offset. For each OLSa list, a speech comprehension threshold (SCT) was computed, which is the difference of the sound pressure of the speech signal and the sound pressure of the noise of the last 20 sentences. For example, an SCT of −7 dB indicates that the participant was able to comprehend 50 % of the sentences that were presented 7 dB below the masking-noise level.

#### 2.2.2. Transcranial alternating current stimulation

tACS was applied using a battery-driven NeuroConn DC Stimulator Plus (NeuroConn GmbH, Ilmenau, Germany). Stimulation electrodes were placed in a bipolar montage on Cz (7 × 5 cm) and bilaterally over the primary auditory cortices on T3 and T4 (4.18 × 4.18 cm) according to the international 10-20 EEG system (Figure 1B). The impedance was kept below 10 kΩ by applying a conductive paste (Ten20, D.O. Weaver, Aurora, CO, USA).

The stimulation signal corresponded to the envelope of the concurrent speech signal (i.e., envelope tACS). As control condition, anodal and cathodal direct current (DC+, DC−) matched to the duration of the speech signal were stimulated as well as no electrical stimulation (sham, see Figure 1B). The envelope-tACS was generated by extracting the envelope of each OLSa sentence. Here, the absolute values of the Hilbert transform of the audio signal were computed and then filtered with a second order Butterworth filter (10 Hz, low-pass). To maximize tACS efficacy, peaks and troughs exceeding 25 % of the highest absolute peak were scaled to 100 %. This way, the peak-to-peak intensity was normalized.

Kubanek and colleagues (2013) reported a lag of around 100 ms of entrained neural oscillations in auditory cortex relative to the entraining speech signal. Here, in order to find the time lag at which auditory stimulation and tACS were aligned in auditory cortex (highest behavioral effect) and also to test how tACS modulates speech comprehension across different time lags, envelope-tACS was assigned to 6 different delay conditions (see Figure 1B; referred to herafter as “time lag”). Envelope tACS was initiated between 0 and 250 ms (50 ms step-size) after the onset of the speech signal. The tACS signal (sampled at 44100 Hz) was delivered via MATLAB to the D/A converter and fed into the external signal input of the DC Stimulator.

#### 2.2.3. Data Analysis

For each participant, we computed the SCT for the six tACS and for the three control conditions (see Figure 2A). First, we wanted to demonstrate that envelope-tACS modulates sentence comprehension. Since we expected that each participant has an individual preferential tACS-time lag, and that sentence comprehension is not explained by specific time lags on the group level, we did not perform an ANOVA on the six time lags. Instead, we selected each participant’s best SCT independent of the time lag and contrasted it with the SCT in the sham condition with a t-test. A significant effect for this contrast is required to assume a modulatory effect of envelope-tACS on sentence comprehension.

**Figure 2.**
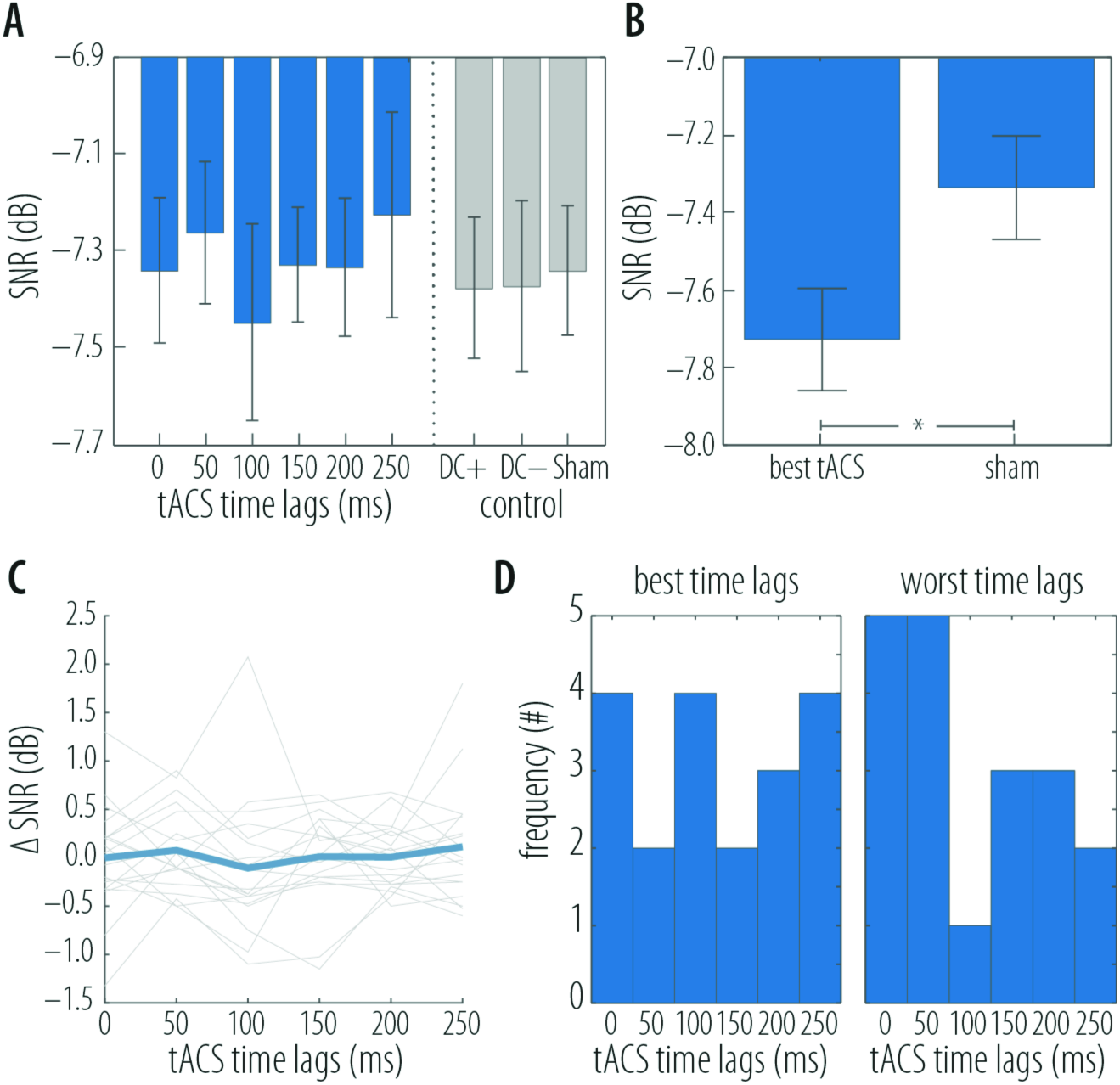
Sentence comprehension thresholds at different time lags. **A**. Bar graphs represent sentence comprehension thresholds (SCTs; signal-to-noise ratio in dB) averaged across all participants at each tACS time lag (blue bars), as well as for the control conditions DC+, DC-, and sham (gray bars). Note that lower values indicate better performance. Error bars display standard error of the mean. **B**. Grand average of lowest SCTs (i.e., best tACS) and sham SCTs. The asterisk indicates the significant difference. Error bars display standard error of the mean. **C**. Δ-SCTs of the six tACS time lags (i.e., tACS-SCT-Sham-SCT). The light gray lines display individual SCTs, the thick blue line displays the mean across individuals. Again, lower values indicate better performance. **D**. Histograms displaying how often a specific time lag resulted in best performance (left panel) and worst performance (right panel).

Next, in order to show that sentence comprehension improves and declines with tACS depending on the time lag we baseline-corrected the SCTs of the six tACS conditions by subtracting the SCT of the sham condition (i.e. Δ-SCT). Here, a negative value indicates improvement in sentence comprehension due to tACS and a positive value indicates a decline in sentence comprehension due to tACS (Figure 2C). We decided to select the sham condition as baseline condition instead of one of the DC conditions for two reasons. First, the SCTs of sham, DC+ and DC− did not differ significantly from each other. Second, as sham is the only condition where no electric stimulation was applied at all, this represents best an inidiviual’s actual sentence comprehension of speech-in-noise.

In order to show that speech comprehension is modulated sinusoidally by tACS along the different time lags, we pursued two, ideally converging approaches: Curve fitting of the behavioral performance over time lags in the time domain, as well as an analysis of spectral power of the performance time series in the frequency domain.

##### Cruve fitting of performance time series

We fitted a sine wave to each participant’s Δ-SCTs, comparable to the approach of Neuling et al. (2012; see also Naue et al., 2011). To show that a sine wave describes the modulation in sentence comprehension best, we also performed a linear fit and a quadratic fit on the Δ-SCTs of each participant. For the sinusoidal fit, the following equation was fitted: ***y = b + a* sin*(*ƒ * 2π* x +c*)**, where *x* corresponded to the tACS time lags [0;250 ms]. The parameters *b, a, ƒ*, and *c* were estimated by the fitting procedure, where *b* was the intercept, *a* was the amplitude of the sine wave, *ƒ* the frequency, and *c* the phase shift. Parameter *b* was bound between −1 and 1, *a* was bound between 0 and 2 (note that the Δ-SCTs were not likely to exceed a value of 1), *ƒ* between 2 and 8 (we expected that a modulation frequency would be in the theta range comparable to the syllable rate of the sentences), and *c* between 0 and 4*π. The initial parameters inserted in the function were randomly drawn from within the range of restrictions of each parameter, respectively. For the linear fit (***y = ax + b***), we estimated the slope *a* and the intercept *b*. The quadratic fit (***y = ax^2^ + bx + c***) was computed to estimate the linear coefficient *b*, the quadratic coefficient *a*, and the intercept *c*. The model fits were computed with the lsqcurvefit function with Matlab (Optimization Toolbox) that allowed for 1000 iterations in order to find the best model. The sinusoidal fit was computed 1000 times for each participant. We then selected the best fit out of these 1000 estimates, in order to avoid premature, suboptimal model fits based on local maxima or minima. The best fit was selected by the lowest Bayesian information criterion (BIC; Schwarz, 1978), a goodness-of-fit parameter based on the log-likelihood of the fit. A lower BIC indicates a better fit.

Next, the BIC was calculated for each fit, in order to compare the goodness-of-fit of the sinusoidal, quadratic, and linear fits. The BIC corrects for an unequal number of parameters and thereby eliminates the advantage of the sinusoidal fit due to a higher number of parameters. BIC scores were compared descriptively. That is, the fit with the lower BICs is the best to describe the modulation of the data.

We evaluated the significance of the sinusoidal fit in itself by performing within-subject permutation tests on each participant’s fit (Ernst, 2004): We took the previously estimated sinusoidal parameters (intercept, amplitude, frequency, and phase), and fitted the permuted x-axis (i.e., time lags) to the sine wave. Six time lags allowed for 720 permutations in total. For each of the permutations the BIC was computed, which resulted in an individual sampling distribution of BICs for each participant. Then, the 5th-percentile BIC of each distribution was determined. If a participant’s empirical BIC from the original sinusoidal fit was below the 5th-percentile BIC of their individual BIC permuted distribution, this sinusoidal fit was classified as significant. For illustration purpose only, permuted BICs were z-transformed and the within-subject sampling distributions were expressed in relative frequencies. The data are thus transformed to a common scale across participants and allows for plotting the average sampling distribution (see Figure 3C).

**Figure 3.**
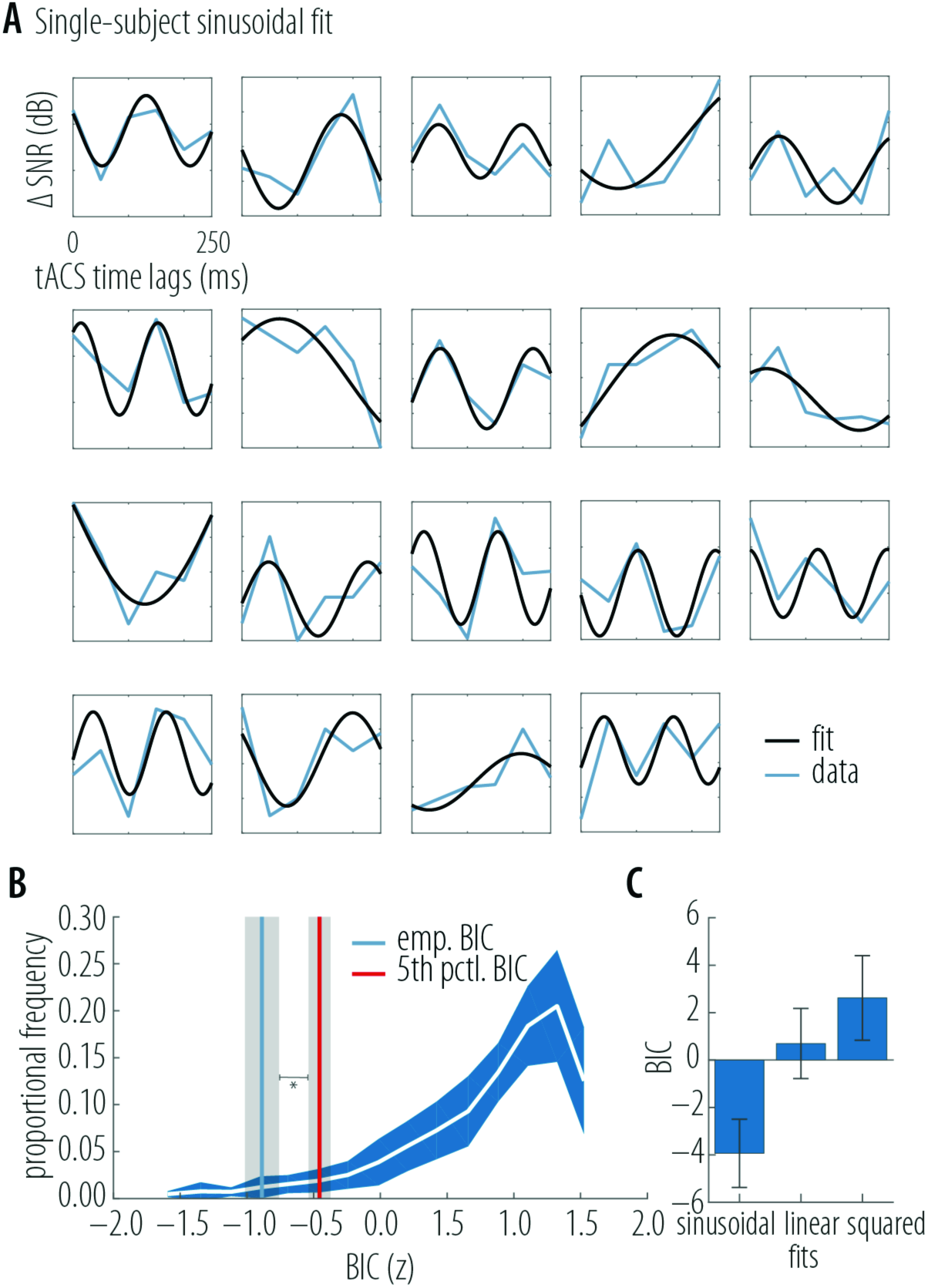
**A**. Single-subject sinusoidal fit. The black curves display the sinusoidal fit. The blue curves are baseline-corrected sentence comprehension thresholds (Δ-SCTs). **B.** Within-subject permutation test. The white line graph displays the grand average of the within-subject sampling distributions of the model-fit Bayes Information criteria (BIC). The within-subject distributions were z-transformed prior to averaging (x-axis). The y-axis indicates the proportional frequency of the BICs. The Blue shade represents the standard error of the mean of the BIC sampling distributions. The red vertical line displays the average BIC (in z-score) of the 5th percentile of each participant. The blue vertical line displays the averaged empirical BIC in z-score. The gray shade of each line, respectively, indicates the standard error of the mean of the empirical BIC and the 5th-percentile BIC. The asterisk marks the significant difference between the percentiles of the empirical BICs and the 5th percentile**. C**. Contrast of goodness-of-fits (i.e., BICs). Bar graphs display the averaged goodness-of-fit (BIC) for each of the three fits. A lower BIC indicates a better fit. Error bars show the standard error of the mean.

##### Analysis of spectral power in the performance time series

To provide corroborating evidence for a cyclic modulation of sentence comprehension across the time lags, we computed the power spectral density of each participant’s Δ-SCTs time series. Prior to the frequency analysis, individual time series of performance were demeaned, interpolated by a factor of 2, multiplied with a hanning window, and tapered to an NFFT length of 320 data points. (Note that interpolation did not qualitatively affect the statistical results.) Individual power spectra were derived as the magnitude of the frequency spectrum.

In order to assess statistical significance of peaks in this spectrum, we derived a permuted null distribution of power spectra by randomly flipping the label of sham versus tACS (for an analogue approach see Fiebelkorn et al., 2013). In brief, in each iteration, the sign of each of the Δ-SCTs in the time domain was flipped randomly to create a new time series prior to frequency analysis. This was performed 5000 times per participant, and the respective power spectra yielded a permuted null distribution of power spectra that were to be expected for random condition assignment.

Each empirically observed mean spectral power value was thus assessed as to whether it exceeded the 95-% percentile of this null distribution. Additionally, a false coverage statement-rate (FCR) correction was applied, correcting for the number of tested frequencies, effectively yielding a one-sided 98.7-% confidence band (Alavash et al., 2017; Benjamini and Yekutieli, 2005).

##### Descriptive analysis of the sentence envelopes

In a last analysis, we aimed to characterize the relationship between the derived peaks in intelligibility modulation and the frequency characteristics of the stimulation sentence envelopes. Therefore, we also estimated the power spectral density on these envelopes. The resulting power spectra were multiplied with the frequency vector to correct for the 1/f shape and to allow for peaks in higher frequencies to become more prominent.

## 3. Results

The aim of the present study was to find out whether envelope-tACS modulates sentence comprehension in noise and to specify the nature of the modulation. Participants listened to sentences with different signal-to-noise ratios. Their ability to comprehend the presented sentences was quantified by the sentence comprehension threshold (SCT), the signal-to-noise ratio at which participants comprehended 50 % of a sentence. tACS was induced either simultaneously or with a varying time lag between 50 and 250 ms. Figure 2A depicts the SCTs for each time lag (i.e., delay between auditory onset and tACS onset), as well as anodal and cathodal tDCS and sham. To test whether tACS at different time lags modulated speech intelligibility at all, we selected the lowest SCTs (best performance) of each participant’s tACS condition and compared it with the SCTs of the sham condition with a t-test (Figure 2B). The t-test showed that the SCTs of the best tACS-performance was significantly better (lower SNR) than the SCTs in the sham condition (t(18) = −4.1, p < 0.0007, Figure 2B). This effect serves as a prerequisite to further investigate whether tACS modulated speech intelligibility.

After selecting the best SCTs, we inspected the distribution of time lags that led to the best and worst performance across participants. The median time lag that led to the best performance was 100 ms and for the worst was 50 ms. Critically, time lags improving or impeding performance were not consistent across participants. The histograms in Figure 2D depict the number of best performances (left panel) and the number of worst performances (right panel) of each time lag. Critically, the 100-ms time lag was the best for three participants (as were time lags 0 ms and 250 ms) but was the worst time lag only for one participant. On a descriptive level, it appears as if the 100-ms time lag might have been more beneficial than other time lags, despite the strong variability across the time lags of best performance.

Next, we wanted to analyze the modulation of sentence comprehension along the different time lags. First, we baseline-corrected the tACS SCTs by subtracting the sham SCTs (i.e., Δ-SCT; Figure 2C). That way the data reflect performance increase and decrease due to envelope-tACS. Then, a sinusoidal fit, a quadratic fit, and a linear fit were performed on the Δ-SCT, in order to understand the nature of the tACS induced modulation of speech comprehension.

The single-subject sinusoidal fits are displayed in Figure 3A. The average goodness-of-fit is BIC = −3.93. Parameters were estimated resulting in this equation, averaged across participants: ***y = 0.023 + 0.386 * sin(5.12 * 2π * x + 0.998)***. Thus, a sinusoidal fit to the performance time series yielded an average fitted modulation frequency of 5.12 Hz.

By comparison, the average goodness-of-fit of the linear fit was BIC = 0.7 with the following coefficient: ***y = 0.12 + 0.001x***. The quadratic fit had an average goodness-of-fit of BIC = 2.62 with coefficients of ***y = 0.034 - 1.276x + 6.194x^2^***. Across participants, the BICs of the sinusoidal fit were lower than the BICs of the quadratic and linear fit (see Figure 3C). That indicates that a sine wave with an average frequency of 5.12 Hz represents the modulation of speech intelligibility by envelope-tACS relatively best.

In order to further assess statistical significance of the sinusoidal fits, we performed within-subject permutation tests (see Methods in *Curve fitting and spectral analysis of performance time series*). It turned out that in 15 out of 19 participants, the BIC of the empirical fit was in the significant tail of their respective BIC permutation distribution (two-sided binomial test p = 0.019). Figure 3B illustrates the grand average of the within-subject BIC sample distributions and the difference between empirical BICs and 5th-percentile BICs. This finding provides additional evidence that envelope-tACS modulates sentence comprehension in a sinusoidal manner.

In the last analysis of the SCTs, we computed the power spectral density of each participant’s Δ-SCTs. Figure 4A displays the interpolated and demeaned Δ-SCTs on which the computation of the power spectra was performed. The thin gray lines show the individual Δ-SCTs time courses, the thick blue line illustrates the mean of all individual time courses. Figure 4B shows the resulting mean power spectrum in blue. Significance was evaluated by contrasting the empirical power spectrum with the power spectra of a permuted null distribution. That is, whether it exceeded the 95-% percentile of this null distribution (solid gray curve in Figure, 4B). Additionally, a false coverage statement-rate (FCR) correction was applied, correcting for the number of tested frequencies, effectively yielding a one-sided 98.7-% confidence band (dashed gray curve in Figure, 4B). A clear significant peak at 5.38 Hz was observable. This effect serves as further evidence of a cyclic modulation of sentence comprehension by envelope-tACS. The solid and dashed black horizontal lines in Figure 4B indicate the frequency range where the empirical mean power spectrum exceeded the respective confidence intervals.

**Figure 4.**
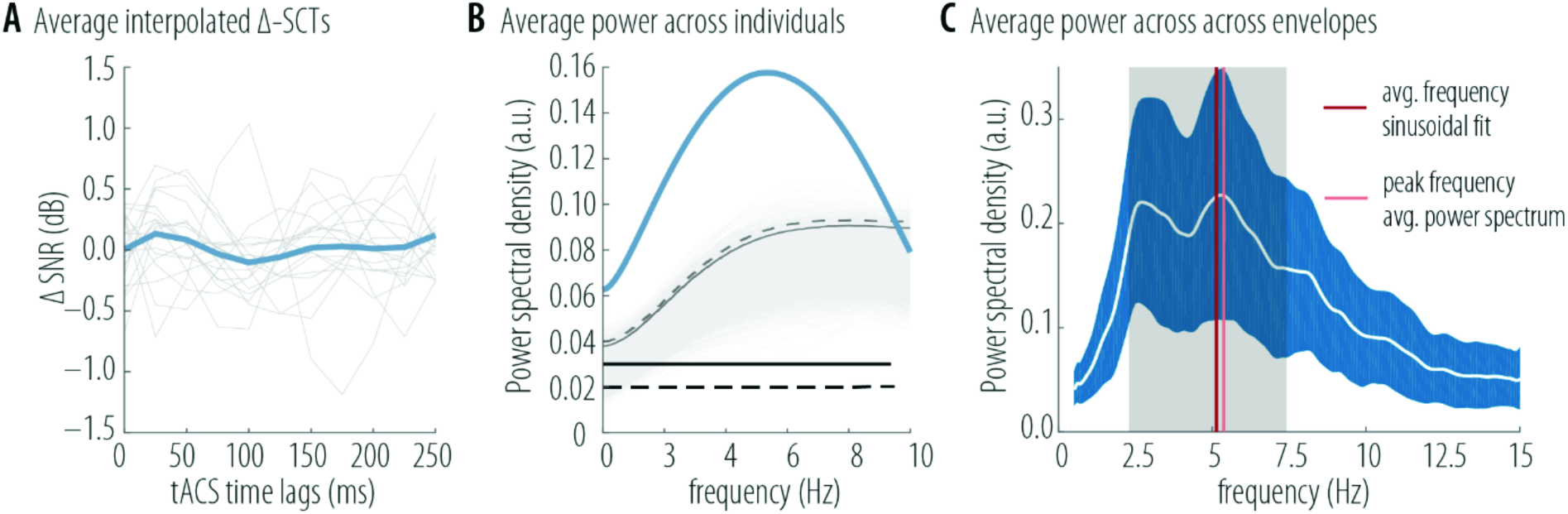
**A**. Δ-SCTs of the six tACS time lags (i.e., tACS-SCT-Sham-SCT). Here, the data are interpolated by the factor 2 and demeaned as preparation for the subsequent computation of the power spectra. The light gray lines display individual SCTs, the thick blue line displays the mean across individuals. Lower values indicate better performance. **B.** Power spectrum averaged across individual power spectra of each participant’s Δ-SCTs (thick blue line). Thin, gray lines display the power spectra of the permuted null distributions. The gray solid curve indicates the 95th percentile confidence interval of the null distribution, the dashed gray curve indicates the FCR-corrected (98.7th percentile) confidence interval of the null distribution. At the frequency bins where the averaged power spectrum exceeds these curves, it is significantly modulated as indicated by the black solid and dotted lines, respectively. **C.** Power spectrum of the sentence envelopes. The white line represents the grand average amplitude spectrum of the speech envelopes. The blue shade reflects 1 standard deviation of all amplitude spectra. The dark red vertical line marks the average frequency of the sinusoidal fit (5.12 Hz). The gray shade indicates the range of the fitted frequencies from the 20th to the 80th percentile. The light red vertical line marks the peak of the average power spectral density (5.38 Hz; see Figure 4C).

Finally, we analyzed the envelopes of the presented sentences that were applied forenvelope-tACS. Figure 4C displays the grand average of the amplitude spectra of the envelopes. Two frequency peaks were most prominent at 2.7 Hz and 5.3 Hz. Importantly, the average frequency of the sinusoidal fits was 5.12 Hz (dark red line) and the peak of the power spectrum was 5.38 (light red line), both close to the 5.3 Hz peak.

## 4. Discussion

The aim of the present study was to show that envelope-tACS in auditory cortex modulates speech intelligibility. We employed a speech intelligibility task in combination with envelope-tACS induced at different time lags relative to the acoustic envelope to identify a delay that would yield the strongest behavioral benefit.

### 4.1. Envelope-tACS modulates sentence comprehension sinusoidally

The present study provides first evidence that sentence comprehension can be modulated by envelope-tACS and that this modulation fluctuates sinusoidally across different time lags. This finding supports previous claims on the important role of cortical entrainment in auditory cortex for speech comprehension (for a review, see Zoefel & VanRullen, 2015). The rationale why envelope-tACS modulates the intelligibility of degraded speech is that the envelope of degraded speech is masked. In the present study, the concomitantly presented noise masked the speech signal. Due to the noise masker, entrainment to the envelope was hampered. tACS is thought to boost the envelope, thereby improving entrainment of the auditory cortex to the speech signal. Similar findings have been reported from a purely acoustic research approach: Koning and Wouters (2012, 2016) enhanced the speech envelope by adding a peak signal at the onsets in each frequency band. With this technique, they could improve intelligibility of degraded speech signals in normal hearing adults and cochlea implant patients.

The sinusoidal fluctuation of speech intelligibility along the different tACS time lags is in line with the results of Neuling et al., 2012. They demonstrated that pure-tone detection depends on the phase of the tACS signal. Entrainment of neural oscillations results in distinct phases of high and low excitability on the population level, thereby generating time windows of a higher firing likelihood (Engel et al., 2001; Lakatos et al., 2005). This synchronization of neural oscillations and external stimuli facilitates the processing of relevant information (Lakatos et al., 2008; Schroeder et al., 2010). Similarly, time lags in the present study varied to assess at which time lag the envelope-tACS signal and the speech envelope are aligned and in phase in auditory cortex. That way, auditory cortex is assumed to synchronize with the tACS signal at the same time it also synchronizes with the speech signal. Consequently, with a different time lag, tACS and the speech signal were not aligned. This misalignment most likely disrupted entrainment of auditory cortex to the speech envelope and disrupted speech comprehension.

However, the sinusoidal modulation of speech comprehension implies that after one cycle of the oscillation or sine wave, tACS modulates sentence comprehension in the same way as it did one cycle before. This is most likely due to the periodicity of entrained neural oscillations and the speech envelope. The circular regularities of the envelope (speech as well as tACS) most probably induced an oscillatory neural response in auditory cortex due to cortical entrainment to the envelope. Consequently, speech comprehension, depending on entrainment to the envelope, oscillates in this circular manner. The average frequency of the fitted sine waves was 5.12 Hz and the peak frequency of the average power spectrum was 5.38 Hz. These almost identical frequencies are within the range of the amplitude modulation of speech at frequencies that are crucial for intelligibility (4 to 16 Hz; Drullman et al., 1994; Shannon et al., 1995). Importantly, these observed frequencies are close to the 5.3-Hz peak of the average amplitude spectrum of the sentence envelopes. That is converging evidence that the effect of envelope-tACS on speech comprehension is indeed related to neural entrainment.

### 4.2. The role of tACS-to-acoustics time lags

The median of the time lags that led to the best sentence comprehension was 100 ms. However, the individual latency of best comprehension performance was found at any of the six time lags ranging from 0 ms to 250 ms. The difference of the frequency of best and worst time lag points out that a time lag of 100 ms tends to result in better sentence comprehension than other time lags. Previous studies used different approaches to analyze temporal phenomena in electrophysiological responses during auditory processing. The most traditional method is the computation of event-related responses, an average of the electrophysiological response across all trials time-locked to the stimulus. In the present study, the median time lag of 100 ms corresponds to the latency of the auditory N100, a negative evoked potential elicited by auditory stimulation after approximately 100 ms (for a review, see Näätänen & Picton, 1987). The N100 has been argued to be a signature of auditory pattern recognition and integration (Näätänen and Winkler, 1999). Although different features of a stimulus can manipulate its characteristics such as amplitude, latency, and origin, the N100 remains one of the robust initial responses to speech (and non-speech sounds) in auditory cortex.

Another method to detect temporal regularities is a cross-correlation between the auditory signal and the respective EEG response. Specifically, for analyzing speech processing, the cross-correlation of the envelope of a sentence and the respective EEG response results in a (spectro-)temporal response function. Here, the peaks of the correlations were found at a latency of 100 ms (Ding and Simon, 2012; Horton et al., 2013). Ding and Simon explain that this latency indicates that the EEG response to a speech signal is driven by the amplitude modulations of the stimulus 100 ms ago. Aiken and Picton (2008) computed transient response models, similar to the cross-correlation of speech signal and EEG response. They reported peaks at a later latency approximately around 180 ms. They also showed that estimated latencies were found within a range from 152 to 208 ms. Although a majority of studies showed that the grand average of cross-correlations converges at a latency around 100 ms, the latencies reported by Aiken and Picton are longer.

Interestingly, an auditory entrainment study by Henry and Obleser (2012) found auditory perception to be modulated by a consistent neural delta phase across participants, whereas the best phase of the entraining stimulus varied across participants. Taken together, these findings lend plausibility to the scenario observed here, namely, that the phase-or time lag between externally entraining stimuli and neural entrainment responses can vary individually, yet that a 100-ms time lag seems optimal in an average sense.

### 4.3. Clinical implications for envelope-tACS

The importance of studying transcranial electric stimulation (tES) lies in its possibilities of clinical applications. Different studies have demonstrated the diverse clinical benefit of tACS (for a summary of therapeutic applications of tES, see Vosskuhl et al., 2015).

The results of the present study indicate that envelope-tACS has the potential of being beneficial for the intelligibility of sentences with a bad signal-to-noise ratio (SNR). People suffering from hearing loss experience bad SNRs constantly. For example, hearing aids are fitted to improve the SNR by filtering, amplifying, and compressing acoustic signals. Digital hearing aids, in contrast to analog hearing aids, even respond to ongoing analyses of the signal and the background on a more complex level (for a review, see Pichora-Fuller & Singh, 2006). In addition to the conventional mode of operation of hearing aids, envelope-tACS could be applied to counteract hearing impairment. Envelope-tACS would then improve the neural SNR in auditory cortex by enhancing cortical entrainment to the relevant speech envelope and thereby improving speech comprehension. However, cortical responses to speech in older adults, the main audience of hearing aid users, need to be investigated, in order to successfully implement envelope tACS in hearing devices.

### 4.4. Limitations

One limitation of this study is that we are unable to draw conclusions about the frequency specificity or the shape of the envelope-tACS effect. It is unclear whether this modulation is solely attained by stimulating the corresponding sentence envelope. In the future, it will be important to test whether sinusoidal tACS in the frequency range of the envelope of the presented sentences will modulate sentence comprehension in a similar or even more effective manner. Since auditory-cortex entrainment to a sine wave will lead to similar fluctuations in cortical excitability, sine-wave tACS might have the same modulatory effect on speech comprehension. To further narrow down the specificity of envelope-tACS, it will also be necessary to apply envelope-tACS of a different sentence. Based on the present findings we would assume that the stimulation of a different envelope would rather cause a disruption of cortical entrainment to the envelope of the presented sentence and hence impede sentence comprehension.

Note, however, that the present data preclude the possibility that the onset of envelope-tACS on its own modulated speech comprehension. If that had been the case, tACS would have induced a phase reset in auditory cortex. It has been previously discussed that the electric current induced by tACS is below the threshold of eliciting action potentials (other than TMS) and does therefore only induce cortical entrainment without a phase reset (Helfrich and Schneider, 2013; Nitsche et al., 2003; Thut et al., 2011).

Another limitation of the study are the sparsely sampled time lags in 50-ms steps within a range of 0–250 ms. Physiologically, 50 ms is a comparably long duration for the temporally fine-grained auditory system. Methodologically, this sampling frequency limits all spectral analyses reported here to frequencies below 10 Hz. Possible faster modulation thus would have not been detectable. In part, the temporally sparse sampling of behavior had technical reasons, since the duration of tACS was limited to 20 minutes per participant per day, it was necessary to limit the number of tACS conditions.

More importantly however, previous studies led us to expect only a modulation of behavioral performance within the frequency range of the entraining stimulus. That has been shown to be the case for auditory detection during tACS entrainment (Neuling et al., 2012; Riecke et al., 2015a) as well as during entrainment to amplitude modulated and/or frequency modulated sounds (Henry et al., 2014; Henry and Obleser, 2012). In our case, we expected behavioral performance to be modulated at or around the main frequency of envelope fluctuation, that is, within the theta frequency range (5 Hz; Figure 4C). Therefore, we did not expect the sinusoidal modulation of speech comprehension to exceed that frequency range.

Lastly, the design of the study as well as the results do not inform us about an actual benefit of envelope-tACS for speech comprehension. Here, further experimentation is needed. For example, if a method is established to determine the individual time lag a priori, envelope-tACS with this time lag should be applied, and sentence comprehension under tACS could be compared to sentence comprehension without tACS.

### 4.5. Conclusions

Altogether, in this study we have demonstrated that envelope-tACS modulates intelligibility of speech in noise, presumably by enhancing and disrupting cortical entrainment to the speech envelope in auditory cortex. This intelligibility modulation itself peaks at or near the peak frequency of the speech envelope. Moreover, the “best time lag” at which tACS appears to be most beneficial varied between participants.

## Acknowledgments

This research was funded by German Research Foundation (Deutsche Forschungsgemeinschaft, DFG Cluster of Excellence 1077 “Hearing4all”).

